# *Cannabis sativa* oxylipin biosynthesis: Genome-wide characterization of lipoxygenase, allene oxide synthase, allene oxide cyclase, hydroperoxide lyase, and 12-oxo-phytodienoic acid reductase gene families

**DOI:** 10.1101/2022.07.30.502131

**Authors:** Eli J. Borrego, Mariah Robertson, James Taylor, Elida Espinoza

## Abstract

*Cannabis sativa* is a global multi-billion-dollar cash crop with numerous industrial uses, including in medicine and recreation where its value is largely owed to the production of pharmacological and psychoactive metabolites known as cannabinoids. Often underappreciated in this role, the lipoxygenase (LOX)-derived green leaf volatiles (GLVs), also known as the scent of cut grass, are the hypothetical origin of hexanoic acid, the initial substrate for cannabinoid biosynthesis. The LOX pathway is best known as the primary source of plant oxylipins, molecules analogous to the eicosanoids from mammalian systems. These molecules are chemically and functionally diverse group of fatty acid-derived signals that govern nearly all biological processes including plant defense and development. The interaction between oxylipin and cannabinoid biosynthetic pathways remains to be explored.

Despite their unique importance in this crop, there has not been a comprehensive investigation focusing on the genes responsible for oxylipin biosynthesis in any *Cannabis* species. This study documents the first genome-wide catalogue of the *Cannabis sativa* oxylipin biosynthetic genes and identified 21 *LOX*, five allene oxide synthases (*AOS*), three allene oxide cyclases (AOC), one hydroperoxide lyase (*HPL*), and five 12-oxo-phytodienoic acid reductases (*OPR*). Gene collinearity analysis found chromosomal regions containing several isoforms maintained across *Cannabis, Arabidopsis,* and tomato. Promoter, expression, weighted co-expression genetic network, and functional enrichment analysis provide evidence of tissue- and cultivar-specific transcription and roles for distinct isoforms in oxylipin and cannabinoid biosynthesis.

This knowledge facilitates future targeted approaches towards *Cannabis* crop improvement and for the manipulation of cannabinoid metabolism.

## 1 Introduction

For millennia, the diploid dioecious shrub *Cannabis sativa*, has been cultivated for its fiber, grain, and pharmacological properties (1). Originating in Asia and now grown globally (2), *C. sativa* is a multibillion-dollar cash crop (3, 4) with many uses (5) including feed (6), textile (7), biofuel (8), medicine (9), and recreation (10, 11). The latter two of which are due to the production of biologically active metabolites, known as cannabinoids, within the cannabis trichomes (12). Cannabinoids are a large chemical family with 130 currently known distinct chemical species (13) grouped into 10 structural types (14). Undeniably, the best understood are the cannabidiol (CBD) (15) and psychoactive Δ^9^-tetrahydrocannabinol (THC) (16). Given their growing therapeutic uses and social acceptance, efforts are underway to produce chemotypes with explicit cannabinoid content (17, 18), platforms for heterologous production (19), and novel cannabinoid analogues (20).

The understanding of cannabinoid biosynthesis has made strides in recent years (12, 19, 21). Briefly, cannabinoids are synthesized from the convergence of a fatty acid and terpene pathway. Hexanoic acid is activated into hexanoyl-CoA by acyl-activating enzyme 1 and subsequently elongated with three molecules of malonyl-CoA by TETRAKETIDE SYNTHASE (also known as OLIVETOL SYNTHASE. TKS functions in concert with OLIVETOLIC ACID CYCLASE to generate olivetolic acid. Geranyl pyrophosphate produced from the non-mevalonate-dependent isoprenoid pathway (MEP) is used by CANNABIGEROLIC ACID SYNTHASE to prenylated olivetolic acid into cannabigerolic acid (CBGA), the first *bona fide* cannabinoid. Cannabigerolic acid can then be shunted into one of at least three oxidocyclization subbranches to produce tetrahydrocannabinolic acid, cannabidiolic acid, and cannabichromenic acid (CBCA) by THCAS SYNTHASE, CBDA-SYNTHASE, and CBCA SYNTHASE, respectively. Later, THCA, CBDA, CBCA may undergo nonenzymatic decarboxylation into the better-known THC, CBD, or cannabichromene. Lastly, cannabinoids may also undergo spontaneous rearrangements that are responsible the structural variation found within this chemical family (12, 13, 22).

Despite the recent advances in the understanding of the cannabinoid biosynthetic pathway, the origin of hexanoic acid (also known as caproic acid) remains unclear. It is often the rate-limiting step in heterogenous systems (20, 23) requiring hexanoic acid as a feedstock (19). In *planta*, trichome-specific transcriptomic analysis observed co-expression of genes involved in oxylipin biosynthesis, namely *LIPOXYGENASE* (*LOX*) and *HYDROPEROXIDE LYASE* (*HPL*), with genes previously established in cannabinoid biosynthesis (24, 25). This has prompted the hypothesis that the oxylipin pathway provides the substrate required for cannabinoid biosynthesis. Further, this putative oxylipin-cannabinoid interaction invites exploration to understand the role of the oxylipins in cannabinoid biosynthesis and *Cannabis* biology.

Oxylipins are analogous to the mammalian eicosanoids, and are a large group of oxidized fatty acid-derivatives possessing potent signaling activities that govern a multitude of physiological processes including, growth, development, and defense (26). In plants, the majority of oxylipin biosynthesis occurs via the LOX pathway. It begins with the regio- and stereo-specific incorporation of molecular oxygen at either the 9^th^ - or 13^th^ -carbon of linoleic (C18:2) or linolenic (C18:3) acid by 9- or 13-LOX isoforms (27). The resulting 9- or 13-oxylipins can be fluxed into at least one of seven sub-branches to produce chemically diverse groups of distinct chemical species, including alcohols, aldehydes, divinyl ethers, esters, epoxides, hydroxides, hydroperoxides, ketols, ketones, and triols. Though the biological role for the vast majority of plant oxylipins remains to be deciphered, investigations of selected members of 13-oxylipins, namely jasmonates and green leaf volatiles (GLVs) have spearheaded a framework towards understanding their physiological and ecological roles.

Jasmonates are best known for their roles in providing defense against insects and necrotrophic pathogens (28). Jasmonates are cyclopentones produced through the ALLENE OXIDE SYNTHASE (AOS) subbranch (29, 30). Following the oxygenation of C18:3 to 13(*S*)-hydroperoxyoctadecatrienoic acid (13-HPOT) by 13-LOX activity, AOS (29, 31, 32) converts 13-HPOT to 12,13(*S*)-epoxy-octadecatrienoic acid (12,13-EOT) and ALLENE OXIDE CYCLASE (AOC) (33) converts 12,13-EOT to (+)-*cis*-12-oxo-phytodienoic acid (12-OPDA), the first jasmonate in the pathway. LOX, AOS, and AOC are closely associated with each other and participate in substrate channeling during jasmonate biosynthesis (34).Aside from serving as a substrate for downstream reactions, 12-OPDA possesses its own distinct activity (35–37). 12-OPDA is reduced via 12-OXO-PHYTODIENOIC ACID REDUCTASE (OPR) (38, 39) to 9*S*,13*S*-OPDA to 3-oxo-2(2′[*Z*]-pentenyl)-cyclopentane-1-octanoic acid (OPC-8:0) and then undergoes three rounds of β-oxidations to produce (+)-7-iso-jasmonic acid (JA) (40). Hexadecatrienoic acid (C16:3) may also serve as substrate for jasmonate biosynthesis, eliminating the need for a round of β-oxidation (41–43). While a multitude of JA derivates have been identified (29), (+)-7-iso-jasmonoyl-L-isoleucine (JA-Ile) is the best explored, owed to its service as a ligand in receptor-mediated perception and subsequent signaling (44).

GLVs are best known as the scent of cut grass and are involved in defense signaling, plant-to-plant and plant-to-insect communication (45). They are C_6_ volatile aldehydes, alcohols, and their acetyl esters produced through the HPL subbranch (46, 47). Here, the 13-hydroperoxy fatty acids produced from 13-LOX activity from either C18:2 or C18:3 are cleaved, respectively, by 13-HPL (48–50) into the hexanal or (3*Z*)-hexenal and 12-oxo-(9*Z*)-dodecenoic acid. The latter of which is the progenitor of the traumatin sub-group, of which some members display signaling activity independent of GLVs (51). Additionally, C16:3 may also serve as a fatty acid substrate for GLV biosynthesis, yielding (7*Z*)-10-oxo-decenoic acid in place of traumatin (52). The GLV aldehydes are reduced to alcohols through reductase (53) and acetylated through acetyltransferase (54). Finally, (3Z)-hexenal can also be isomerized through (3Z):(2E)-ENAL ISOMERASE (55, 56).

Thus, this study sought to elucidate the *Cannabis* oxylipin biosynthetic pathway as a source for cannabinoid substrate and molecular signals important in defense and development. Here, a census was conducted on the *C. sativa* genome to catalog the major oxylipin biosynthetic genes from the LOX, AOS, AOC, and OPR gene families. Respectively, their chromosomal locations, conserved protein domains, genetic structures, and phylogenetic relationships were determined. Collinearity analysis identified several genes in syntenic regions with *Arabidopsis* and tomato. The promoter analysis revealed evidence for tissue- and stimulus-specific transcriptional regulation, which stands in agreement with expression observed from a *Cannabis* transcriptome atlas and publicly-available transcriptomes. Finally, gene co-expressional network and functional enrichment analysis identified genetic modules that implicate oxylipin biosynthetic genes in distinct *Cannabis* physiological processes.

## 2 Materials and Methods

### Gene model identification, protein domain analysis, and subcellular localization prediction

The *Cannabis sativa* representative genome [cs10](https://www.ncbi.nlm.nih.gov/assembly/GCF_900626175.2/) (57) was surveyed for LOX, AOS, HPL, AOC, and OPR gene models using the NCBI BLASTP algorithm against the *Arabidopsis* sequences as the input queries. The corresponding public transcriptome data was explored through the NCBI Genome Data Viewer (58) to obtain gene models and evidence of transcription and putative translation. Amino acid sequences were examined against the NCBI Conserved Domain Database for the presence of canonical domains using [Batch CD-Search](https://www.ncbi.nlm.nih.gov/Structure/bwrpsb/bwrpsb.cgi) (59, 60). Conserved peptide sequence motifs were determined using [MEME 5.4.1](https://meme-suite.org/meme/tools/meme) (61, 62) for up to 25 motifs and any number of repetitions. Subcellular localization prediction analysis was performed using [DeepLoc 1.0](https://services.healthtech.dtu.dk/service.php?DeepLoc-1.0) (63), [LOCALIZER](https://localizer.csiro.au/) (64), Plant-mSubP (65), and [TargetP 2.0](https://services.healthtech.dtu.dk/service.php?TargetP-2.0) (66) through their web servers with default settings.

### Multiple sequence alignment, phylogenetic analysis, and genetic structure visualization

Peptide sequences were aligned through the online version of [MAFFT](https://mafft.cbrc.jp/alignment/server/index.html) (67, 68) using the FFT-NS-I iterative refinement strategy, leaving gappy regions, and with other settings as default. Phylogenetic trees were generated through the MAFFT website using neighbor-joining, Jones-Taylor-Thornton substitution model, with estimated heterogeneity. Trees were tested with bootstrapping of 1000 resampling and drawn with [FigTree 1.4.1](http://tree.bio.ed.ac.uk/software/figtree/). Gene structures were visualized with [TBtools v1.098689](https://github.com/CJ-Chen/TBtools) (69) using the NCBI *Cannabis sativa* Annotation Release 100.

### Promoter analysis of cis-regulatory elements

Putative cis-acting regulatory elements were determined using the 1.5 kb upstream nucleotide sequences of gene models, analyzed with [PlantCARE](http://bioinformatics.psb.ugent.be/webtools/plantcare/html/) (70), and promoter diagrams were constructed with TBtools.

### Comparative genomic analysis

Gene collinearity between *C. sativa*-*A. thaliana* (reference genome [TAIR10.1](https://www.ncbi.nlm.nih.gov/genome/4?genome_assembly_id=380024)) and *C. sativa*-*S. lycopersicum* (representative genome [SL3.0](https://www.ncbi.nlm.nih.gov/genome/7?genome_assembly_id=393272)) was performed using MCScanX (71) feature of TBtools with an e-value 1e-10 and 5 BLAST hit cutoffs.

### Transcriptomic profiling, regulatory network analysis

RNA-Seq SRP achieved data was retrieved from GEO Datasets, SRP234963 (72) and SRP168446 (73), and trimmed with fastp v0.23.1 (74) using default settings. Transcripts Per Million (TPM) were calculated using [salmon](https://salmon.readthedocs.io/en/latest/salmon.html) (75) mapping-based mode against a decoy-aware transcriptome file. Corrections were made for fragment-level GC content and random hexamer primer biases. Transcriptome indices were computed from the cs10 reference transcriptome against its respective genome.

Gene co-expression networks were generated through an iterative weighted correlation network analysis (WGCNA) using [iterativeWGCNA](https://github.com/cstoeckert/iterativeWGCNA) with default parameters (76, 77). Relationships between highly interconnected gene modules were explored through an eigengene score correlation matrix visualized through [Cytoscape](https://cytoscape.org/) (78).

Functional enrichment analysis was performed using [g:Profiler](https://biit.cs.ut.ee/gprofiler/gost) (79) using orthologues identified through [PLAZA 5.0](https://bioinformatics.psb.ugent.be/plaza/) (80) and cross-referenced against the [Kyoto Encyclopedia of Genes and Genomes](https://www.genome.jp/kegg/) (81–83) and [Gene Ontology Database](http://geneontology.org/) (84, 85).

## 3 Results

### Cannabis oxylipin biosynthetic genes

To establish a foundation for *Cannabis* oxylipin biology, the *C. sativa* oxylipin biosynthetic genes were determined. The *C. sativa* representative genome was queried against selected *Arabidopsis* LOX, AOS, AOC, and OPR amino (19)acid sequences with BLASTP to detect candidate *C. sativa* orthologues. The analysis identified 21 LOX, six CYP74, three AOC, and five OPR distinct gene models (Table 1).

**Table 1.** Genes encoding LOX, CYP74, AOC, and OPR gene families in C. sativa and their predicted peptide sublocalization

| Annotation | Locus | Chromosome | Location | Sense | mRNA | Protein | AA | Subcellular localization analysis |  |  |  |
| --- | --- | --- | --- | --- | --- | --- | --- | --- | --- | --- | --- |
|  |  |  |  |  |  |  |  | DeepLoc | LOCALIZER | mSubP | TargetP |
| CsLOX1 | LOC115719608 | 2 | 48717953..48726423 | - | XM_030648706.1 | XP_030504566 | 860 | Cytoplasm | - | Cyto | - |
| CsLOX2 | LOC115718693 | 2 | 48399032..48404403 | - | XM_030647516.1 | XP_030503376 | 955 | Plastid | - | Plastid | - |
| CsLOX3 | LOC115724062 | 9 | 45035547..45040076 | - | XM_030653517.1 | XP_030509377 | 872 | Cytoplasm | - | Cyto | - |
| CsLOX4 | LOC115709296 | 3 | 1987993..1993483 | + | XM_030637370.1 | XP_030493230 | 859 | Cytoplasm | - | Cyto | - |
| CsLOX5 | LOC115720291 | 2 | 48437789..48443653 | - | XM_030649445.1 | XP_030505305 | 848 | Cytoplasm | - | Cyto | - |
| CsLOX6 | LOC115721132 | 2 | 48616833..48624053 | - | XM_030650384.1 | XP_030506244 | 859 | Cytoplasm | - | Cyto | - |
| CsLOX7 | LOC115719336 | 2 | 48633009..48641095 | - | XM_030648324.1 | XP_030504184 | 871 | Cytoplasm | - | Cyto | - |
| CsLOX8 | LOC115722276 | 9 | 58835655..58849094 | + | XM_030651442.1 | XP_030507302 | 868 | Cytoplasm | - | Cyto | - |
| CsLOX9 | LOC115721268 | 2 | 3929913..3935080 | + | XM_030650533.1 | XP_030506393 | 875 | Cytoplasm | - | Cyto | - |
| CsLOX10 | LOC115722275 | 9 | 58804514..58809886 | + | XM_030651441.1 | XP_030507301 | 868 | Cytoplasm | - | Cyto | - |
| CsLOX11 | LOC115723988 | 9 | 58775266..58790926 | + | XM_030653445.1 | XP_030509305 | 873 | Cytoplasm | - | Cyto | - |
| CsLOX12 | LOC115718785 | 2 | 181580..185609 | - | XM_030647611.1 | XP_030503471 | 838 | Mitochondrion | Mitochondrion | Endoplasm | mTP |
| CsLOX13 | LOC115712696 | 4 | 8039511..8043316 | + | XM_030641024.1 | XP_030496884 | 935 | Plastid | cTP | Plastid | cTP |
| CsLOX14 | LOC115707105 | 1 | 89971772..89976562 | - | XM_030634953.1 | XP_030490813 | 931 | Plastid | cTP | Plastid | - |
| CsLOX15 | LOC115719612 | 2 | 93914675..93922215 | + | XM_030648714.1 | XP_030504574 | 929 | Plastid | cTP | Cyto | - |
| CsLOX16 | LOC115719614 | 2 | 93928923..93935412 | + | XM_030648717.1 | XP_030504577 | 926 | Plastid | - | Cyto | - |
| CsLOX17 | LOC115720530 | 2 | 93947032..93955004 | + | XM_030649675.1 | XP_030505535 | 1293 | Plastid | - | Plastid | - |
| CsLOX18 | LOC115719613 | 2 | 93802540..93809737 | + | XM_030648716.1 | XP_030504576 | 929 | Plastid | cTP | Plastid | cTP |
| CsLOX19 | LOC115719615 | 2 | 93892217..93901839 | + | XM_030648718.1 | XP_030504578 | 922 | Plastid | cTP | Plastid | - |
| CsLOX20 | LOC115719616 | 2 | 93836391..93848870 | + | XM_030648719.1 | XP_030504579 | 917 | Plastid | - | Plastid | - |
| CsLOX21 | LOC115719617 | 2 | 93822415..93829745 | + | XM_030648721.1 | XP_030504581 | 909 | Plastid | cTP | Plastid | - |
| CsAOS1 | LOC115723125 | 9 | 2271373..2273482 | + | XM_030652553.1 | XP_030508413 | 525 | Plastid | cTP | Plastid | cTP |
| CsAOS2 | LOC115703182 | X | 6627388..6629208 | + | XM_030630703.1 | XP_030486563 | 482 | Peroxisome | - | Celmemb | - |
| CsAOS3 | LOC115708244 | 1 | 22613465..22615178 | + | XM_030636464.1 | XP_030492324 | 483 | Plastid | - | Celmemb | - |
| CsAOS4 | LOC115722144 | 9 | 55962020..55963631 | + | XM_030651275.1 | XP_030507135 | 503 | Plastid | - | Celmemb | - |
| CsAOS5 | LOC115703186 | X | 6682344..6684188 | - | XM_030630707.1 | XP_030486567 | 503 | Plastid | - | Celmemb | - |
| CsHPL | LOC115698766 | 8 | 51951025..51958560 | + | XM_030625939.1 | XP_030481799 | 485 | Plastid | - | Plastid | - |
| CsAOC1 | LOC115701324 | 8 | 54784220..54785234 | + | XM_030629083.1 | XP_030484943 | 183 | Cytoplasm | - | Celmemb | - |
| CsAOC2 | LOC115700991 | 8 | 54774852..54776132 | - | XM_030628671.1 | XP_030484531 | 249 | Plastid | cTP | Plastid | cTP |
| CsAOC3 | LOC115699327 | 8 | 35236892..35238108 | + | XM_030626678.1 | XP_030482538 | 257 | Plastid | cTP | Plastid | cTP |
| CsOPR1 | LOC115706391 | 1 | 27999501..28002834 | - | XM_030634031.1 | XP_030489891 | 452 | Peroxisome | cTP | Plastid | - |
| CsOPR2 | LOC115706393 | 1 | 27991920..27994962 | - | XM_030634033.1 | XP_030489893 | 392 | Peroxisome | - | - | - |
| CsOPR3 | LOC115706394 | 1 | 27977420..27994954 | - | XM_030634034.1 | XP_030489894 | 392 | Peroxisome | - | Cyto | - |
| CsOPR4 | LOC115703964 | 1 | 27889405..27891439 | - | XM_030631189.1 | XP_030487049 | 392 | Peroxisome | - | Cyto | - |
| CsOPR5 | LOC115722904 | 9 | 35363370..35365841 | - | XM_030652270.1 | XP_030508130 | 371 | Peroxisome | cTP | Cyto | cTP |

The genes were found asymmetrically distributed both across and within the 10 *C. sativa* chromosomes (Fig 1) with the largest concentration of genes was found on Chromosome 1, 2, and 9. Over 60% of the LOX isoforms were found on Chromosome 2, and with the exception of two, were in large gene clusters. *CsLOX1*, *2*, *5*, *6*, and *7* were located in a 327 Kb region in the center of Chromosome 2, while *CsLOX15*, *16*, *17*, *18*, *19*, *20*, and *21* were near the outer arm in a 152 Kb region (S1 Fig.). A smaller 74 Kb region on Chromosome 9 contained the cluster of *CsLOX8*, *10*, and *11* towards the end of a chromosomal arm. *CsLOX9* and *12* were relatively close together on Chromosome 2, while Chromosomes 1, 3, and 4 contained single LOX isoforms, *CsLOX14*, *4*, and *13*, respectively.

**Fig 1.**
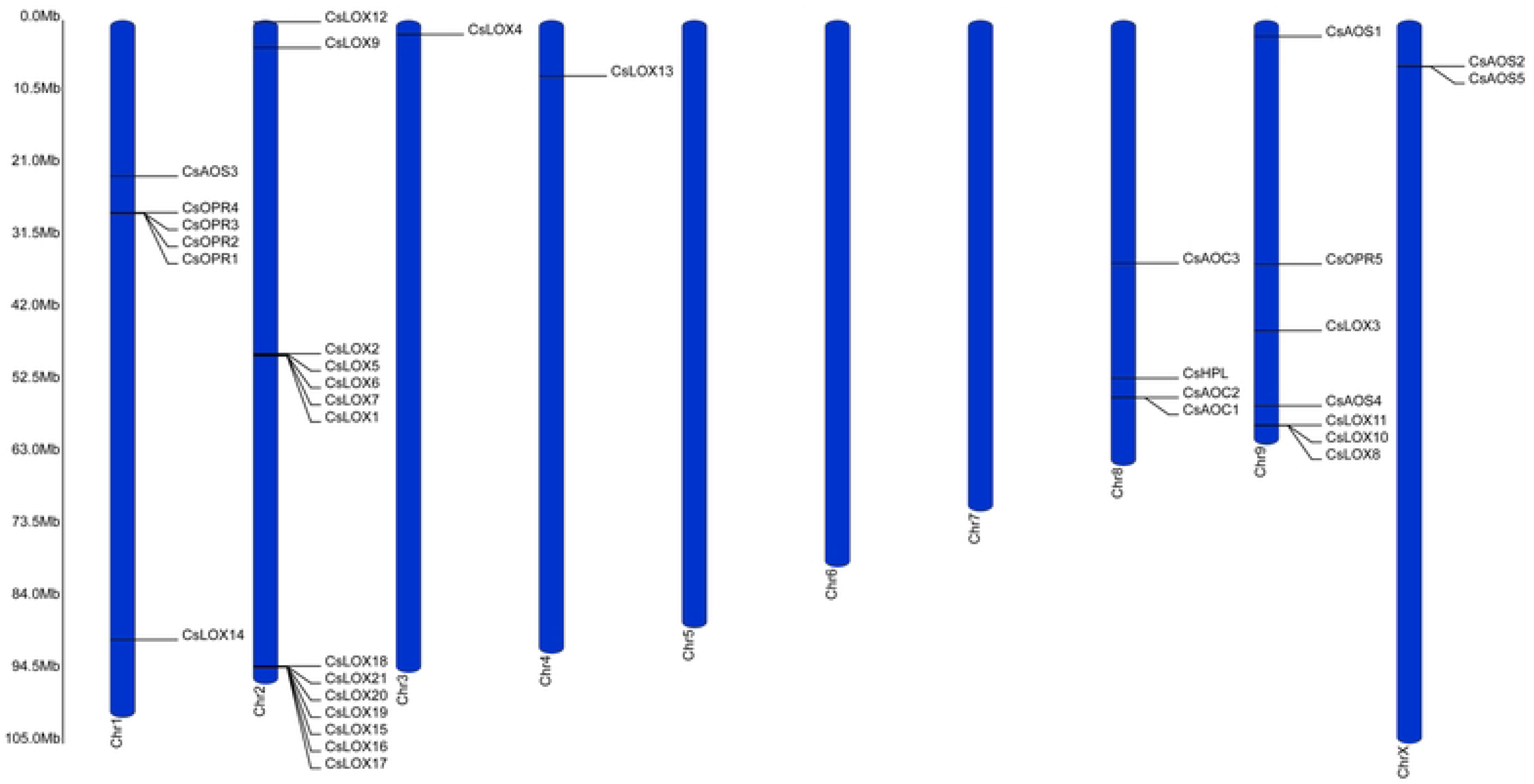
Distribution of the LOX, AOS, HPL, AOC, and OPR gene families across the *C. sativa* genome. Ruler depicts chromosome length.

Similar to the *LOX* genes, members of the CYP74, AOC, and OPR gene families were found in gene clusters at nearly the same proportions. On Chromosome 8, CsAOS2 and 5 were within 57 Kb from each other (S2 Fig.). The other *CYP74* genes, *CsAOS1, CsHPL, CsAOS1,* and *CsAOS4* were on Chromosomes 1, 8, and 9, respectively. *AOC* genes were found exclusively on Chromosome X with *CsAOC1* and *2* only 10 Kb apart (S2 Fig.). Strikingly, with the exception of *CsOPR5* located on Chromosome 9, the entire OPR gene family was confined to a 113 Kb segment on Chromosome 1.

### The C. sativa LOX gene family contains 21 members

To verify that candidate LOX gene models encode functional isoforms, their peptide sequences were examined for the presence of the lipid-associated PLAT and catalytic LOX domains archetypical of LOX proteins (27). Amino acids were examined through the NCBI Conserved Domain Database function and all 21 gene models encoded peptides that displayed both domains, providing support that *C. sativa* possesses 21, *bona fide*, LOX isoforms (Fig. 2A). Unexpectedly, the analysis revealed that *CsLOX17* also contained a RVT2 and a RNAse H-like domain which may indicate the insertion of a retrotransposon in its upstream region (86, 87).

**Fig 2.**
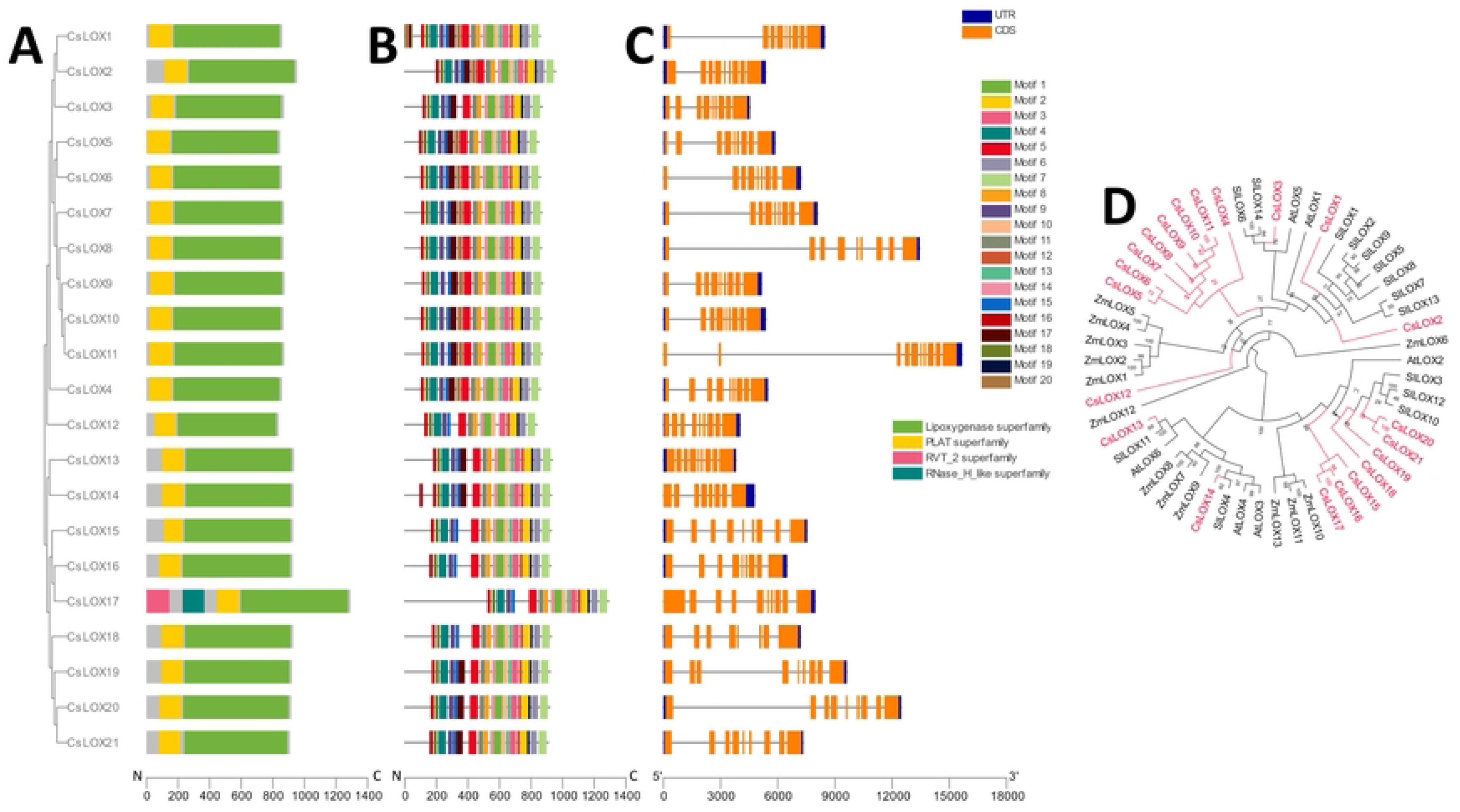
Phylogenetic, genetic, motif, and domain analysis of the CsLOX gene family. **(A)** Cladogram of peptide sequences and conserved domains. **(B)** Distribution of conserved peptide sequence motifs. Colors are described in legend; x-axis represents length of peptides in amino acids. **(C)** Diagram of genetic structure. Blue bars, orange bars, and gray lines represent untranslated regions, exons, and introns, respectively; x-axis represents length of gene in nucleotides. **(D)** Polar cladogram depicting evolutionary relationship with gene families of selected species. Node labels show confidence values from 1000 bootstrap replications.

Analysis of conserved motifs showed that the majority of the LOX isoforms contain similar composition and distribution of amino acid sequence patterns (Fig. 2B). Nearly all isoforms displayed the same 13 motif arrangement, typical of LOX proteins, on the N-terminus, a highly conserved region which requires stringent composition for enzymatic activity. The exception was CsLOX12 which was absent of motif 13. Aside from CsLOX1 and 14, the arrangement of the first 5 motifs on the C-terminus was broadly consistent across all the proteins. However, several isoforms possessed regions where no motifs were detected. In LOX proteins, these regions typically contain highly divergent, transient peptide sequences that direct subcellular localization (88). In support of this notion, their amino acid sequences were tested against four subcellular prediction algorithms, DeepLOC, LOCALIZER, mSUBP, and TargetP (Table 1). A consensus of plastid localization was reached for only CsLOX13 and CsLOX18, along with an agreement of the majority of the software localizing CsLOX2, CsLOX14, CsLOX15, CsLOX17, CsLOX18, CsLOX19, CsLOX20, and CsLOX21 to this organelle. CsLOX1 and CsLOX3-11 were predicted to be found in the cytoplasm. Interestingly, CsLOX12 was predicted to be associated with the mitochondria.

Motifs 17 and 20 located within the LOX domain showed the largest variability across the LOX isoforms. The former was absent from CsLOX3, 13, and 14, the latter was missing from CsLOX19 and 20, and both were missing from CsLOX12, 15, 16, 17, and 18, prompting the speculation of divergent catalytic activity among even closely related members (89).

The *C. sativa* LOX gene models ranged in size from around 4 to 16 Kb and over 75% possessed eight introns (Fig. 2C). A recurrent observation in the genetic structures of some *LOX* family members was the presence of a large first or second intron, e.g., CsLOX1, 6, 7, 8, 11, and 20 comprising of about 50 to 60% of the entire gene. No data is yet available for alternative splicing or transcript variants of these genes to provide insight into the role of these features.

Plant LOXs are typically grouped as 9- or 13-LOXs according to their major enzymatic product and isoforms with similar activity display higher sequence similarity and evolutionary relationships (27). To designate the classification of *C. sativa* LOXs, their peptide sequences were compared to dicot and monocot plant species with well-characterized LOX protein families, namely, *A. thaliana, O. sativa, S. lycopersicum, and Z. mays* (Fig. 2D). CsLOX1, 2, and 3 grouped with the *Arabidopsis* 9-LOXs, AtLOX1 and 5, respectively. Remarkably, eight *Cannabis* LOXs peptides, CsLOX4-11, formed a *Cannabis* specific clade nested within monocot and dicot 9-LOXs. CsLOX12 was difficult to place within the tree, a result which was likely related to the absence of the three variable peptide motifs in its catalytic domain (Fix. 2B). CsLOX13 and 14 clustered with clades containing dicot 13-LOXs. Interestingly, seven *Cannabis* LOX peptides, CsLOX15-21 grouped with the 13-LOX clade, and several of which may represent *Cannabis*-specific 13-LOXs. Thus, *C. sativa* possess eleven 9-LOXs (CsLOX1 – 11), nine 13-LOXs (CsLOX13 – 21), and one that remains to be empirically determined (CsLOX12).

### The *C. sativa* CYP74 gene family contains six members

AOS and HPL belong to the atypical CYP74 clade of the P450 family involved in hydroperoxyl fatty acid rearrangement or dismutation (90). Amino acid sequence analysis found a single CYP74 domain in all six *Cannabis* candidate CYP74s and four distinct patterns of conserved sequence motifs (Fig. 3A, B). The pattern variations were most pronounced towards the N-termini where each possessed a distinctive starting motif. The N-terminus of CYP74s typically possess transient peptide signals involved in directing their subcellular localizations towards plastids, and while subsequent analysis prediction supported this notion in CsHPL and CsAOS1, a consensus among the four software prediction tools could not be reached for CsAOS2-5 (Table 1). For gene structure analysis, CsHPL was notable for the presence of two introns (Fig. 3C), an unusual feature of CYP74 family members, one of which comprised the majority of its gene model sequence.

**Fig 3.**
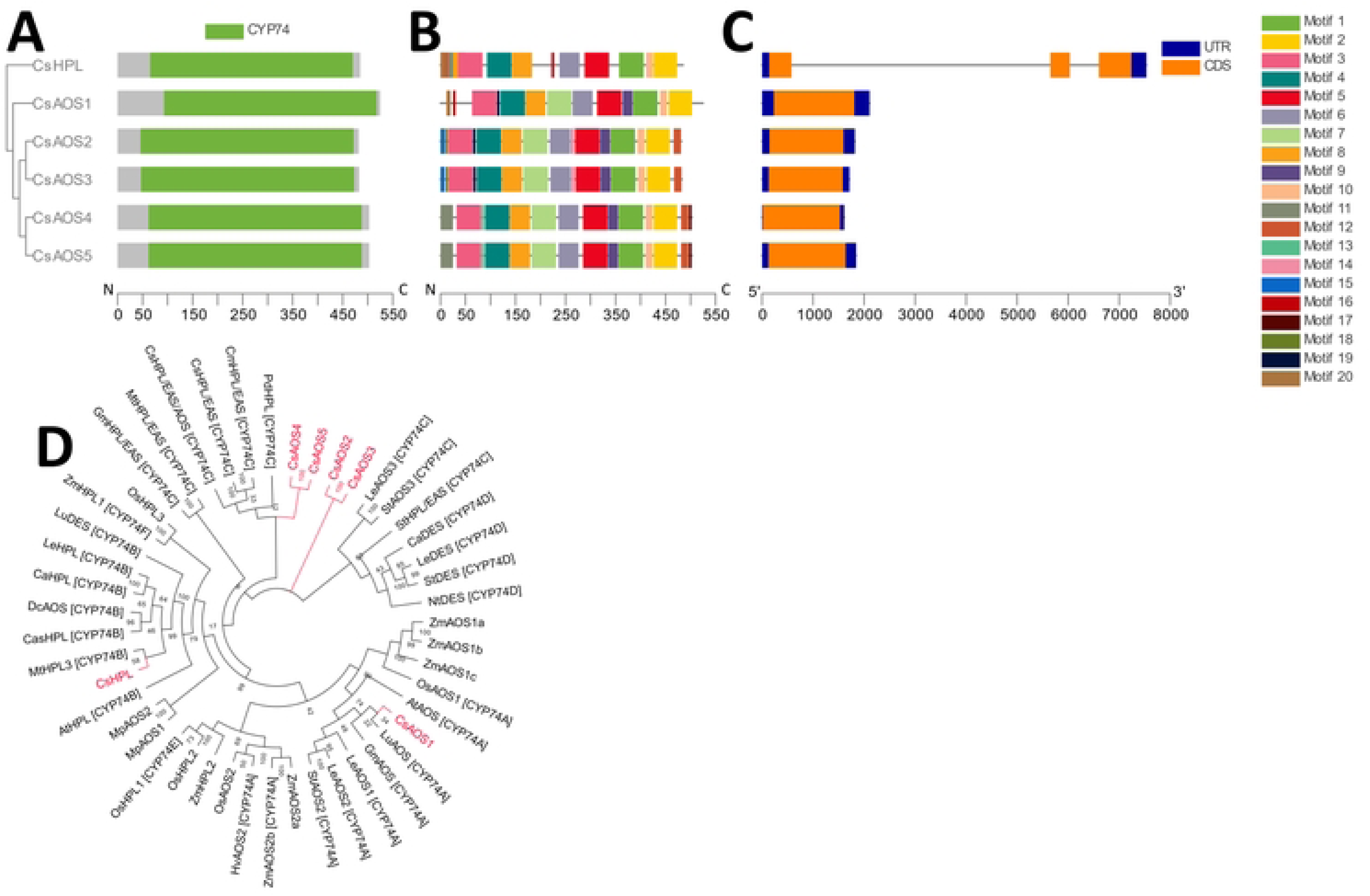
Phylogenetic, genetic, motif, and domain analysis of the *C. sativa* CYP74 gene family. **(A)** Cladogram of peptide sequences and conserved domains. **(B)** Distribution of conserved peptide sequence motifs. Colors are described in legend; x-axis represents length of peptides in amino acids. **(C)** Diagram of genetic structure. Blue bars, orange bars, and gray lines represent untranslated regions, exons, and introns, respectively; x-axis represents length of gene in nucleotides. **(D)** Polar cladogram depicting evolutionary relationship with gene families of selected species. Node labels show confidence values from 1000 bootstrap replications.

While many CYP74 isoforms display multifunctionality (91), catalyzing varying proportions of AOS, EAS, DES, and HPL products, the CYP74A and CYP74B clade members show predominantly AOS and HPL activities, respectively. To understand the potential function of *C. sativa* CYP74 family members, a phylogenetic analysis assessed their relationship within the relatively well-characterized CYP74 subclades from diverse plant species (Fig. 3D). CsAOS1 and CsHPL were grouped within the CYP74A and CYP74B clades, respectively. However, the placement of CsAOS2-4 proved challenging. Nonetheless, CsAOS2 and 3 and CsAOS4 and 5 grouped as pairs roughly within CYP74C.

Taken together, this analysis suggests that *C. sativa* contains at least two dedicated CYP74 enzymes that, given their phylogenies and subcellular localization, are specialized for 13-AOS and HPL activity, thus capable of providing substrate for JA or GLV biosynthesis. Cannabis also contains four CYP74 isoforms with unassigned activities.

### The *C. sativa* AOC gene family contains three members

AOC provides steric hindrance for the stereospecific cyclization of allene oxide to 12-OPDA, the parent jasmonate species. All three CsAOC isoforms contain an AOC domain and similar amino acid sequence patterns on the last six motifs of their C-termini (Fig. 4A, B). CsAOC2 and CsAOC3 displayed substantial variability in predicted motifs at their N-termini, likely corresponding to putative plastid transient peptide sequences (Table 1). CsAOC1 was predicted to localize outside of the plastids. The intron-exon distribution of this gene family differed mildly with CsAOC1 and 2 having three exons while CsAOC3 contained one (Fig. 4C). To understand the evolutionary relationship of the *C. sativa* AOC gene family, their peptide sequences were analyzed phylogenetically with AOC members from other plant species (92). CsAOC1 and 2 clustered close to each other within a dicot specific clade and CsAOC3 grouped into a separate dicot-specific clade that contained all *Arabidopsis* AOCs (Fig. 4D).

**Fig 4.**
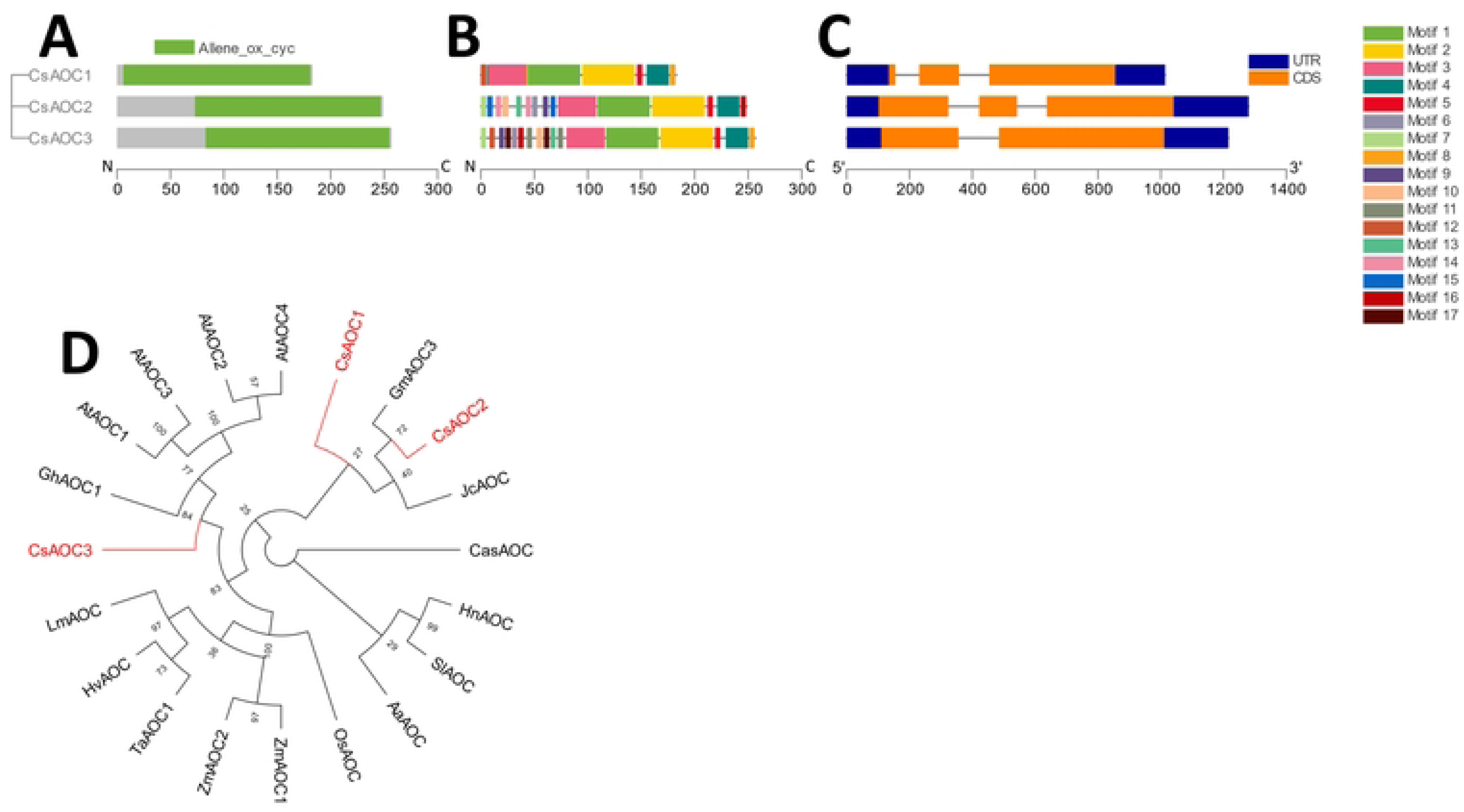
Phylogenetic, genetic, motif, and domain analysis of the CsAOC gene family. **(A)** Cladogram of peptide sequences and conserved domains. **(B)** Distribution of conserved peptide sequence motifs. Colors are described in legend; x-axis represents length of peptides in amino acids. **(C)** Diagram of genetic structure. Blue bars, orange bars, and gray lines represent untranslated regions, exons, and introns, respectively; x-axis represents length of gene in nucleotides. **(D)** Polar cladogram depicting evolutionary relationship with gene families of selected species. Node labels show confidence values from 1000 bootstrap replications.

### The *C. sativa* OPR gene family contains five members

All five identified *C. sativa* OPR candidates were found to contain the conserved Old Yellow Enzyme-like domain necessary for their activity (Fig. 5A) (93). CsOPR1-4 had nearly identical patterns of amino acid sequence motifs and were largely similar to the motif arrangement of CsOPR5, however, the latter isoform lacked four conserved motifs (Fig. 5B). In regard to their genetic structure, a substantial first intron was identified in CsOPR3, resulting in a gene model roughly three times larger than the other isoforms (Fig. 5C). It is important to note that the current gene model of CsOPR3 appears chimeric with CsOPR2 in the current *C. sativa* representative genome (S2 Fig.).

**Fig 5.**
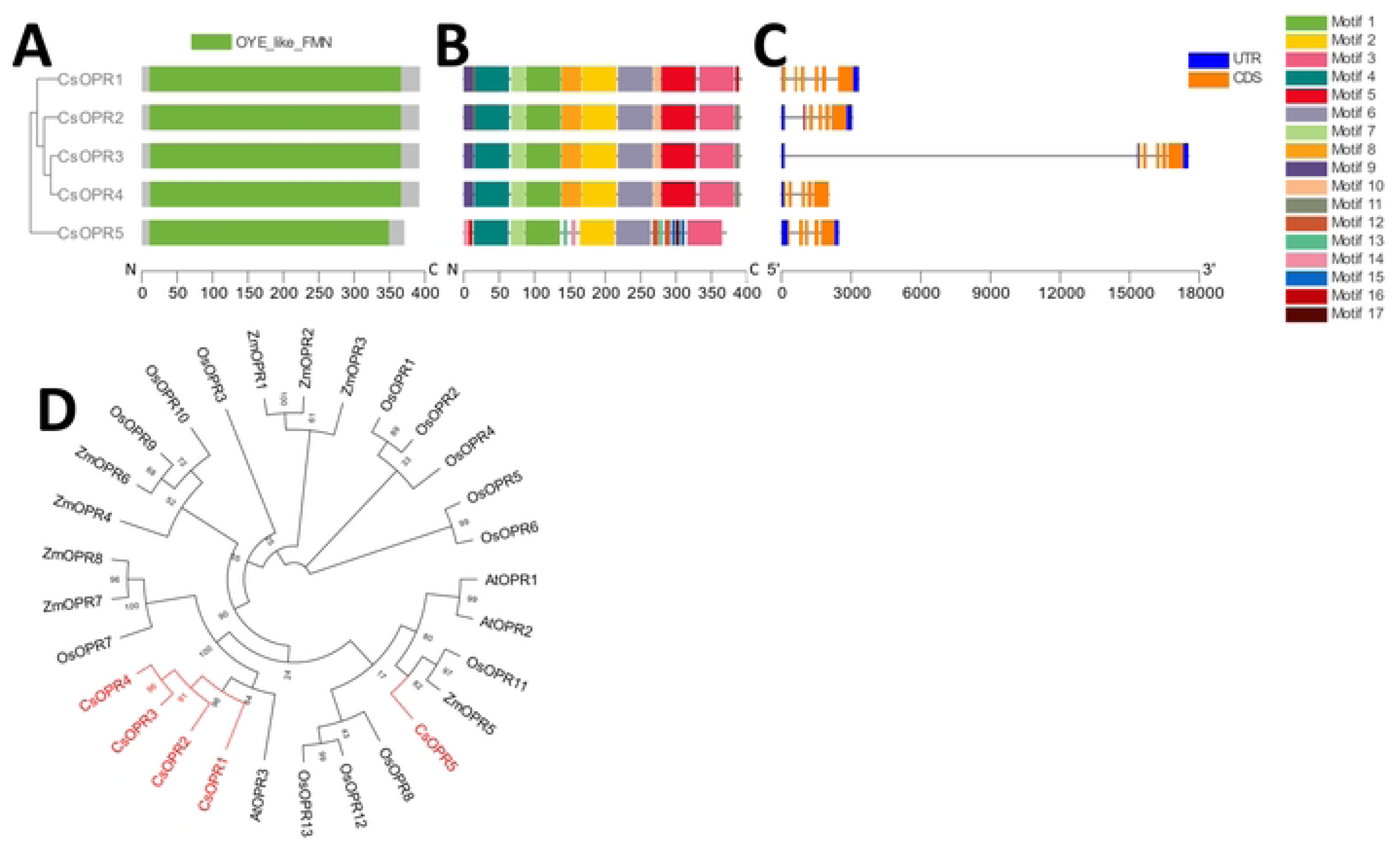
Phylogenetic, genetic, motif, and domain analysis of the CsOPR gene family. **(A)** Cladogram of peptide sequences and conserved domains. **(B)** Distribution of conserved peptide sequence motifs. Colors are described in legend; x-axis represents length of peptides in amino acids. **(C)** Diagram of genetic structure. Blue bars, orange bars, and gray lines represent untranslated regions, exons, and introns, respectively; x-axis represents length of gene in nucleotides. **(D)** Polar cladogram depicting evolutionary relationship with gene families of selected species. Node labels show confidence values from 1000 bootstrap replications.

To investigate the evolutionary relationship between the *C. sativa* OPR gene family members, the amino acid sequences were compared to OPR family members from dicots and monocots. CsOPR1-4 formed a phylogenetic clade with the JA-producing AtOPR3. CsOPR5, however, was grouped with Type-II OPRs (Fig. 5D). Taken together, *C. sativa* contains four Type-I OPR isoforms that are likely involved in classical JA biosynthesis.

### Some LOX, CYP74, and AOC members are syntenic across *Cannabis*, *Arabidopsis*, and tomato

To infer the ancestry of the *C. sativa* oxylipin pathway, a genetic collinearity analysis was performed to compare physical co-localization of oxylipin biosynthetic genes in the genomes of *C. sativa* against two dicot genomes with well characterized oxylipin biosynthetic pathways, *A. thaliana*, and *S. lycopersicum* (tomato) (Fig. 6). Remarkably, three LOX genes displayed collinearity across all three species. Of the 9-LOXs, *CsLOX2* was in a syntenic region with *AtLOX1* and *SlLOX5 (*also known as TOM*LOXE*). Two 13-LOXs display collinearity, *CsLOX14* with both *AtLOX3* and *AtLOX4* and with *SlLOX4/ LOXD,* while *CsLOX13* was collinear with *AtLOX6* and *SlLOX11.* Two *C. sativa* CYP74 genes were in syntenic regions with tomato genes. *CsAOS2* matched with *SlDES* (also known as *LeDES*) and *CsHPL* matched with both *SlHPL* and *SlAOS3.* Two *CsAOC* genes were collinear with either plant species although with dissimilarities. *CsAOC2* was in a syntenic gene block with *SlAOC* and *CsAOC3* matched with both *AtAOC1* and *AtAOC4*.

**Fig 6.**
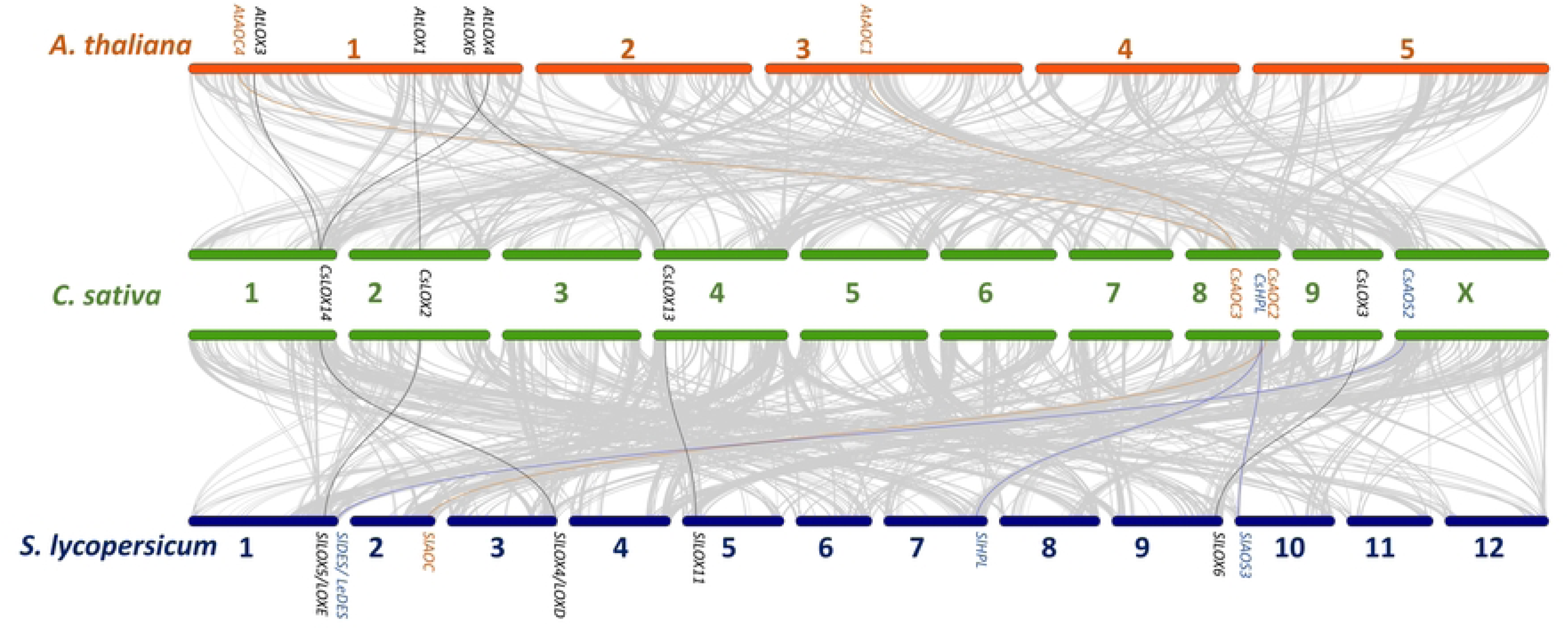
Oxylipin biosynthetic gene collinearity between *A. thaliana*, *C. sativa*, and *S. lycopersicum*. The grey lines represent collinear genetic blocks between the genomes of *A. thaliana* and *C. sativa* and between *C. sativa* and *S. lycospersicum*. Bars represent chromosomes and the black, blue, and orange lines represent the collinear pairs of LOX, CYP74, and AOC gene families, respectively.

### Promoters of *C. sativa* oxylipin biosynthetic genes carry tissue-specific, conditional, and phytohormone-inducible cis-acting regulatory elements (CAREs)

To elucidate the involvement of *Cannabis* oxylipin genes in diverse developmental and stress responses, an *in silico* promoter analysis was conducted to identify cis-regulatory elements involved in plant tissue-specific expression (endosperm, meristem, mesophyll, seed), adaptation to environmental conditions (anaerobic, circadian rhythm, drought, light, and low temperature), and phytohormone signaling (ABA, GA, IAA, JA, and SA).

All promoters varied considerably in the content, arrangement, and position of their CAREs (Fig. 7 and S1 Table). Overall, the oxylipin biosynthetic gene promoters possessed CARE motifs with lengths of 8 to 32 nucleotides. For the LOX gene family, the promoter of *CsLOX12* gene was found to contain the most regulatory elements with 32 and followed by *CsLOX16* and *21*, each with 28, while *CsLOX2, 7, 11*, and *17* displayed the least, with around 10 each. The *Cannabis*-specific 9-LOX group (CsLOX4-11) displayed the most motifs followed by the amplified GLV-producing 13-LOXs (CsLOX15-21). With the exception of CsLOX5, all contained several regulatory elements involved in conditional responses, though few contained elements involved in tissue expression. CAREs involved in light responses dominated most promoters with all genes having at least three such motifs. Interestingly, CsLOX12 light-responsive motifs constituted over 60% of all its identified motifs. Promoters associated with the presence of phytohormone inducible cis-regulatory elements varies considerably, spanning from 0 to 8 regulatory elements. With the exception of CsLOX2, 4, 7, 8, and 17, all had at least one CARE related to ABA responses. Incidentally, CsLOX2, 4, 7, 8, and 17 were also among the few members that showed motifs involved in GA responses, suggesting a role mediated by ABA-GA antagonism (94). SA-responsive motifs were limited and predominately associated with the 9-LOXs.

**Fig 7.**
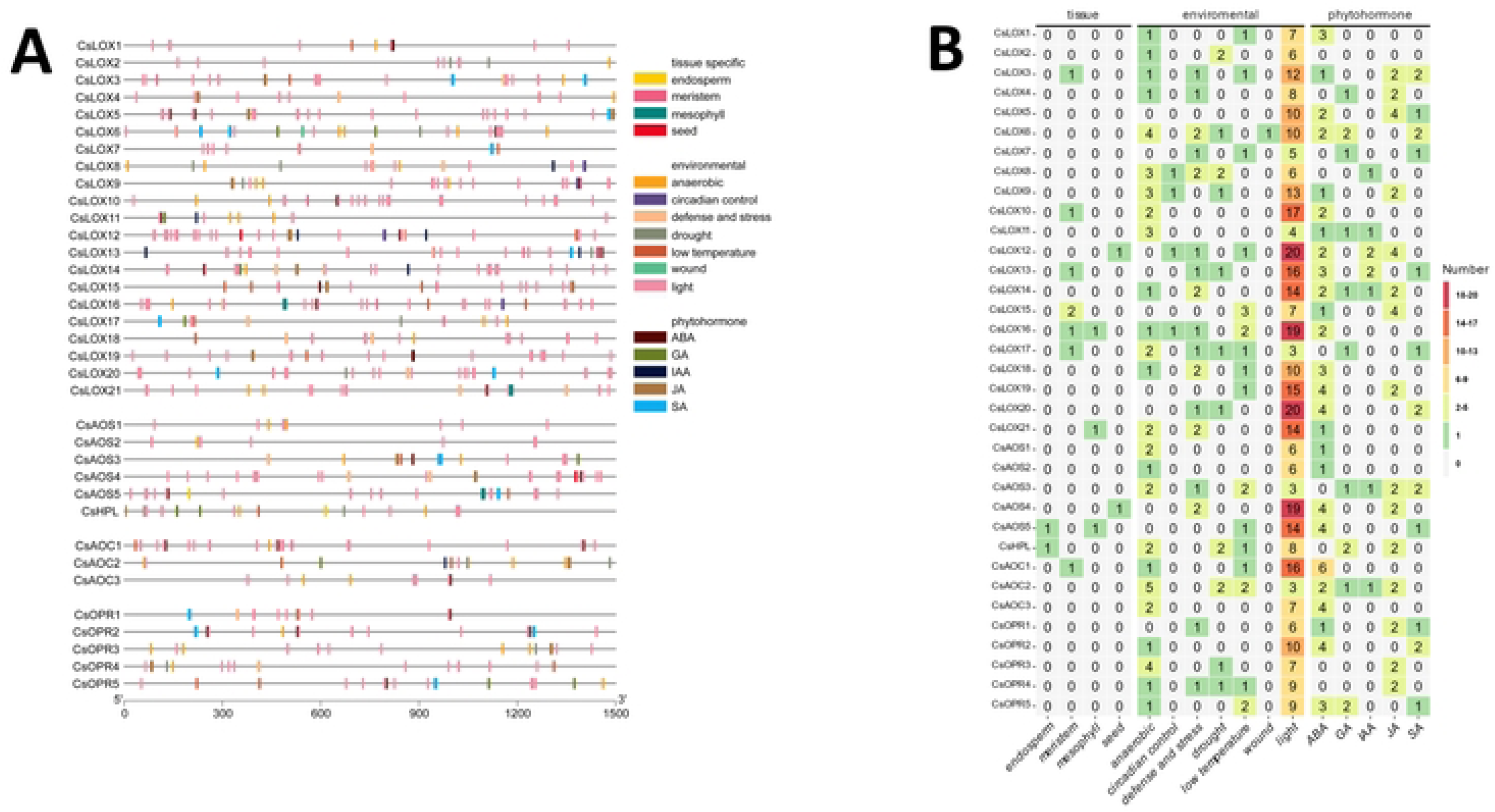
*Cis*-acting regulatory element distribution across *C. sativa* oxylipin biosynthetic gene promoters. (**A**) Physical distribution of motifs throughout a 1.5 kb region upstream of corresponding gene model. Gene symbols are presented on the left with grey line representing promoter regions with motif positions. Legend describes motifs associate with tissue-specific expression, environmental responses, and phytohormone inducibility. (**B**) Summary of *cis-*acting regulatory motif content. Heatmap depicts number of motifs identified in gene promoter region. The y-axis depicts *C. sativa* oxylipin biosynthetic genes and x-axis depicts associated condition for motif. Cells values are number of motifs identified and are colored according to legend.

Of the CYP74 gene family, the largest number of CAREs were identified on promoters of CsAOS4 and 5, followed by CsHPL (Fig. 7). Relatively few were found on the putative JA-producing CsAOS1, namely those associated with anaerobic conditions, light, and ABA signaling responses. Unlike the pattern observed for LOXs, the disparity between light-related motifs and other CAREs was not as dramatic in the CYP74 family, with the exception of the CsAOS4 and 5 pair. Nearly all CYP74s had ABA-responsive motifs, with the exception of CsAOS3 and CsHPL which were also the only ones to possess GA-related motifs. CsAOS3, 4, and CsHPL each had two JA-related motifs, while CsAOS1 had none; this suggests a limited contribution of positive feedforward control of JA biosynthesis.

Analysis of the AOC gene family found only a single tissue-type related to CAREs across the promoters of all members: CsAOC1 for meristem expression. While non-light conditional-response motifs were the most numerous in CsAOC2 compared to the other two isomers, the opposite was seen for light-responsive motifs, where only three were found in CsAOC2. This was a sharp contrast with CsAOC1, which showed more than twice the number of these CAREs compared to its paralogues (Fig. 7). No AOC was found to have SA-related, and only CsAOC2 had JA-related motifs, while all had ABA-related motifs.

Promoters of the OPR gene family contained similar patterns of CAREs compared with the other gene families, albeit with reduced CARE content. No tissue-related CARE was detected for any OPR member. However, all other members possessed motifs for anaerobic responses, with the exception of CsOPR1 (Fig. 7). ABA motifs were found in CsOPR1, CsOPR2, and CsOPR5. CsOPR5 was also the only member to show GA motifs, and JA motifs were found in CsOPR1, 3, and 4. The notable difference in CARE patterns found within the JA-producing OPRs (CsOPR1-4), suggests that the 12-OPDA to JA balance is maintained in *Cannabis* through transcriptional control of distinct OPR members responses to the plant’s environment and during phytohormone signaling (95).

### *C. sativa* oxylipin biosynthetic genes show tissue-, developmental-, and cultivar-specific expression

Promoter analysis suggested that expression of *C. sativa* oxylipin biosynthetic genes would display distinct patterns. To understand the transcriptional profile of oxylipin biosynthetic genes and to elucidate their role in *Cannabis* development and physiology, the gene expression of 23 tissues from a *C. sativa* transcriptome atlas (72) were examined (S2 Table). Overall, the major contributor associated with levels of expression appeared to be the identity of the specific gene family member, i.e., patterns of expression levels were generally consistent across the tissues examined (Fig. 8A). Moreover, several members in all gene families had few if any detected transcripts across all tissues, suggesting a limited role for those isoforms under basal conditions or in the tissues profiled.

**Fig 8.**
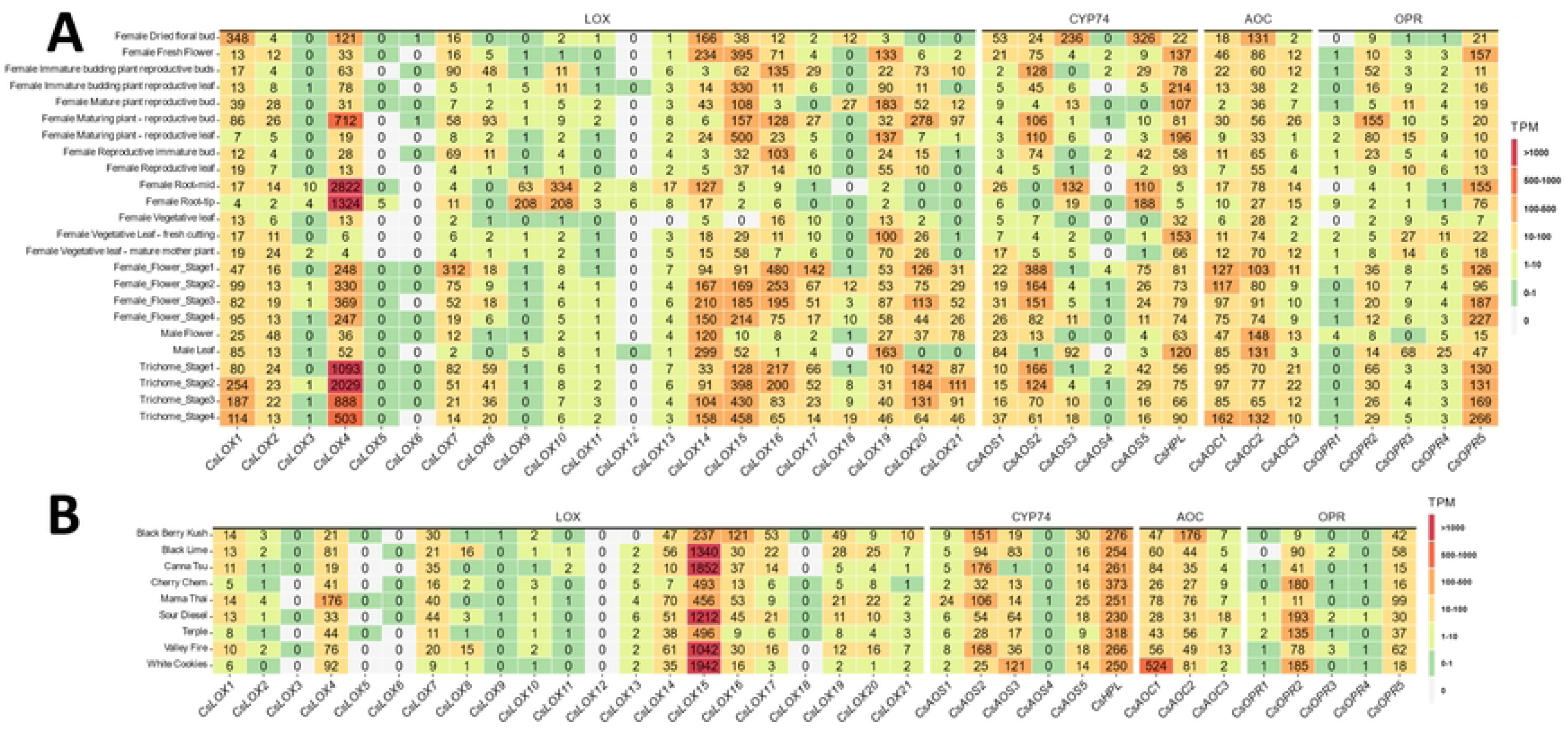
Heatmaps showing expression of oxylipin biosynthetic genes from the C. sativa gene expression atlas (A) or female trichomes of diverse marijuana lines (B). Cells values are rounded TPM values and are colored according to legend

Of the 9-LOXs, *CsLOX1, 2, 4, 7* and *8* showed pronounced levels of expression across most tissues. CsLOX4 is particularly notable as the highest expressed oxylipin biosynthetic gene analyzed, by one order of magnitude, in root and trichome tissues (Fig. 8A), highlighting the probable importance of this isoform grouping within the *Cannabis*-specific 9-LOX clade (Fig. 2). Curiously, *CsLOX9* and *10* had minimal levels of expression in all but root tissues, where their levels were 20-fold higher compared to other tissues. With respect to the 13-LOXs, with the exception of *CsLOX13* and *18*, all had elevated expression in reproductive-related tissue. However, under these basal conditions, only *CsLOX14* showed modest expression in roots. Remarkably, despite possessing the greatest number of identified CAREs (Fig. 7B), nearly no expression was detected for *CsLOX12* (Fig 8A) which implies a non-basal role for this gene.

In regards to the CYP74-related genes, the likely non-JA producing *CsAOS2* displayed the most abundant expression levels, predominantly in early stages of flower and trichome development (Fig 8A, S3). The JA-producing, *CsAOS1*, also showed its greatest levels of expression in the female flowers, while minimally expressed in other tissues. While *CsAOS5* showed modest levels of expression across tissues, transcripts of its closest paralogue *CsAOS4* were only detected at low levels in few tissues. For the GLV- and cannabinoid-producing *CsHPL,* expression remained consistent across tissues, disregarding low levels in root tissues (Fig. 8A). Interestingly, *CsHPL* expression remained relatively consistent across all stages of flower and trichome development (S3 Fig.).

In the majority of tissues, *CsAOC2* showed the most uniform levels of expression from this gene family (Fig. 8), suggesting this member is the prominent isoform involved in JA biosynthesis under during basal conditions. *CsAOC1* and *CsAOC3* showed moderate to modest levels of expression across most tissue types with several exceptions, prompting the idea that these members participate in inducible processes.

Only *CsOPR2 and 5,* displayed mentionable levels of expression. *CsOPR2* transcript levels were an order of magnitude higher than the other Type II paralogues, suggesting this is the major JA-producing isoform under basal conditions (Fig. 8). It is also notable that the majority of its expression was in tissues related to flower structures. The sole Type I OPR, *CsOPR5* displayed even greater levels of expression in flower tissues, especially developing trichomes, raising the possibility for the involvement of CsOPR5 during reduction or detoxification of metabolites during trichome development.

To understand the variation in gene expression across *Cannabis* diversity, transcriptomes of trichomes from nine cultivars of mixed ancestry (73) were profiled for their expression of oxylipin biosynthetic genes(S3 Table). With the exception of expression of *CsLOX15,* of which transcript levels showed nearly a bimodal distribution across the cultivar tested, i.e., Black Berry Kush, Cherry Chem, Mamma Thai, and Terple possessed half of the expression compared to the five other lines (Fig. 8B). Few differences were observed in the other oxylipin biosynthetic gene expression. In particular, decreased expression was seen in *CsLOX14, CsLOX16, and CsOPR2* from Cherry Cham, Terple, and Black Berry Kush, respectively, relative to the other *Cannabis* lines. On the other hand, Mama Thai and Black Berry Kush showed elevated levels for transcripts of *CsLOX4* and *CsAOC1,* respectively, and trichomes of White Cookies had increased expression of both *CsAOS3* and *CsAOC1.* Interestingly, in sharp contrast to expression levels seen in the transcriptome atlas, transcripts of *CsLOX15* dominated in the trichomes of these cultivars followed by those of *CsLOX4*. In agreement with the oxylipin-derived hexanoic acid hypothesis (96), the expression of *CsHPL* levels were also two- to 10-fold higher compared to the other CYP74s, likely owed to the selection of these cultivars for high cannabinoid production.

### *Cannabis* oxylipin biosynthetic genes are found in gene networks associated with stress responses, growth, and development

To assign molecular and cellular functions to the *Cannabis* oxylipin biosynthetic genes, an iterative weighted gene co-expression network analysis was performed on a *Cannabis* transcriptome atlas (72) to identify clusters of genes with highly correlated expression. To inform their roles in physiological processes, the resulting gene modules were analyzed for functional term enrichment. In total, 23 modules containing at least one *Cannabis* oxylipin biosynthetic gene were identified with seven modules containing multiple genes (Fig. 9A). The four largest modules contained between 1130-1852 genes and comprised nearly 30% of the current *C. sativa* gene models; one module contained *CsAOS5*, while the other three modules contained multiple oxylipin biosynthetic genes: *CsLOX3* and *10*, *CsOPR3* and *4*, and *CsLOX5*, *9* and *CsOPR1*. The 15 smallest modules had under 200 genes each and together accounted for only 5% of the *C. sativa* transcriptome.

**Fig 9.**
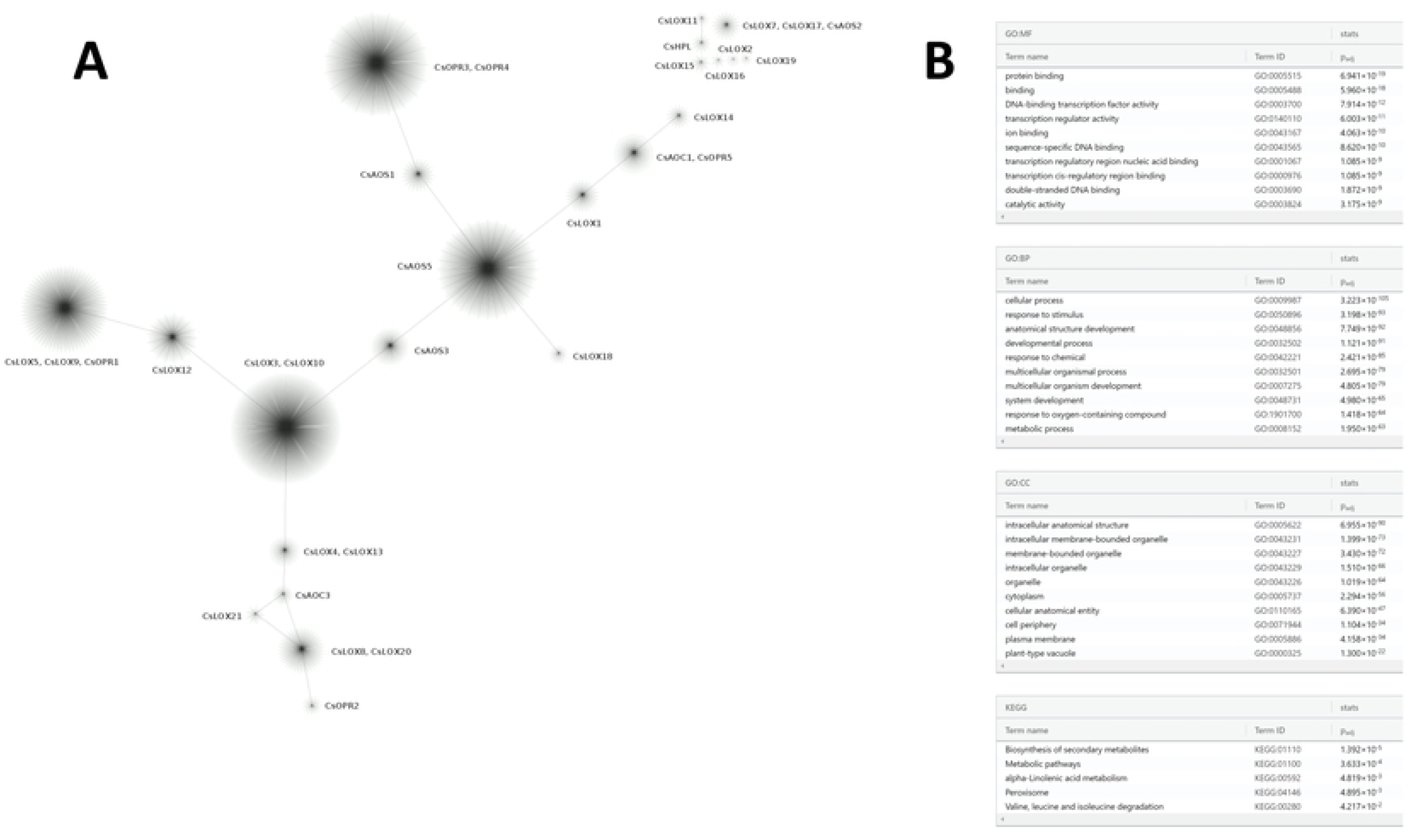
Weighted co-expression genetic network derived from the *C. sativa* transcriptome expression atlas. **(A)** Clusters represent modules of highly connected genes containing at least one oxylipin biosynthetic gene. **(B)** Top ten terms for functional enrichment analysis of gene modules containing oxylipin biosynthetic genes: Gene Ontologies for Molecular Functions (MF), Biological Processes (BP), and Cellular Components (CC) and KEGG.

Functional enrichment analysis of the entire oxylipin gene-associated co-expression network for Gene Ontology and KEGG identified terms related to molecular functions, biological process, and cellular component orthologs (Fig. 9B). Among the top 10 terms for molecular functions, 60% were related to nucleic acid binding and transcription. Of the biological processes, 80% were related to development or physiological responses. For cellular components, 40% and 30% described organelle- and membrane-related terms, respectively. KEGG terms described secondary metabolism, particularly alpha-linolenic acid metabolism and branch-chain amino acid degradation. As expected, individual modules were enriched for differential functional terms (S4 Table). Taken together, these results suggest that even closely related *Cannabis* oxylipin biosynthetic genes are specialized for discrete functions.

## 4 Discussion

Enthusiasm is growing to understand the physiological and ecological functions of plant oxylipins (90). JA has long served (97) as the model oxylipin, and over the last fifty years, many aspects of its biosynthesis, metabolism, signaling, and activities have started to be defined (98–101). However, this knowledge has been generated predominantly in the model plant species, *Arabidopsis*, and investigations into oxylipin biology in crop species have been launched only in recent years. Here, we cataloged the major oxylipin biosynthetic genes families of the agro-economically important crop species, *C. sativa,* described their expression patterns, and delineated connections between their putative physiological functions.

### LOX gene family

This study adds Cannabaceae to the collection of plant families with described LOX gene families, a group which so far includes Aracae: duckweed (102); Actinidiaceae: kiwi (103); Brassicaceae: *Arabidopsis thaliana* (104–106), radish (107), and turnip (108); Caricaceae: papaya (106); Cucurbitaceae: cucumber (109, 110), melon (111), and watermelon (112); Fabaceae: *Medicago truncatula,* peanut (113), and soybean (114); Malvaceae: cotton (115) Muscacea: banana (116); Poaceae: maize (117), rice (105), *Setaria italica* (118), sorghum (119); Polygonaceae: buckwheat (120); Rhamnaceae: Jujubie (121); Rosaceae: apple (122), peach (123), pear (124); Salicaceae: poplar (125); Solanaceae: pepper (126) and tomato (127, 128); Theaceae: *Camellia sinensis* (129); and Vitaceae: grape (130).

Of the 21 *C. sativa* LOX isoforms identified (Fig. 1), 11 are likely responsible for the production of 9-oxylipins, nine are producers of 13-oxylipins, and one member (CsLOX12) remains to be empirically determined. Of the hallmark 9-LOXs AtLOX1 and AtLOX5, two and one *C. sativa* LOX members grouped with each respectively, while eight formed their own clade. Within 9-LOX phylogenies, monocots typically display their own grouping (102, 116, 118), while the divergence of 9-LOXs in dicots is less common; yet still observed in Cucurbitaceae (111), Salicaceae (125), and Vitaceae (130). Interestingly, seven members were identified in nearly one large tandem gene array (S1 Fig.), which fell within the clade containing both monocot and dicot isoforms required for GLV biosynthesis (45, 131). Similar substantial gene amplification of this LOX clade has been found to various degrees, in Cucurbitaceae (109–112), Salicaceae (125), and Vitaceae (130). As *AtLOX2* was not found to be syntenic across the Rosids (106), the gene duplications likely occurred independently in ancestors following the divergence of Rosales, Cucurbitales, Malpighiales, and Vitales.

Proteomic analysis (132) has verified that several lipoxygenases are found with spatial specificity in *Cannabis* flowers and components of their glandular trichomes. Of particular interest was the presence of isoforms from the tandem duplicated 13-LOXs, namely, CsLOX15, 16 and 17 within the trichome head. It is tempting to speculate that these gene duplications may serve to maintain the pool of hexanoic acid substrate available for cannabinoid biosynthesis. In *Cannabis*, tandem gene arrays have previously been implicated in regulating lipid biosynthesis, particularly the ratios of volatile terpenes that subsequently determines the scent of specific cultivars (133). This amplification of the 13-LOX paralogues in a tandem gene array may be due to the origin of the representative *C. sativa* genome (57) from CBDRx, a high CBD producing line. The ubiquity of tandem gene arrays among other *Cannabis* species and cultivars remains to be observed, especially in regards to the agronomic characteristics that drive the selection of economically important cultivars. Interestingly, the 9-LOXs, CsLOX4 and 7 were also found within the flower and trichomes where they are likely to contribute to the 9-oxylipin volatiles of *Cannabis* (134).

### *CYP74* gene family

Despite its importance in oxylipin metabolism, few studies have examined CYP74 gene families from a genome-wide perspective (135–137). In *C. sativa*, six CYP74 isoforms were identified Specifically CsAOS1 and CsHPL, were distinctly found within the JA- and GLV-producing CYP74 clades, respectively (Fig. 3D). Both contained a transient peptide signal sequence predicting localization in plastids (Table 1), although the unique motifs found at their N-terminus suggest an association with different sub-plastid locations. This is similar to tomato where LeAOS was found with the inner chloroplast envelope while LeHPL was targeted to the outer plastid envelope (138). Four isoforms were found with the CYP74C subfamily, a more irregular group of enzymes with members possessing varying levels of AOS, EAS, and/or HPL activity (139). It is interesting to note that CsAOS2 was syntenic to the DES of tomato (Fig. 6), however, a comprehensive survey and validation of *Cannabis* oxylipins remains to be performed to understand if this species indeed produces divinyl ethers. Ultimately, these four isoforms will require biochemical characterization, as even a single amino acid change can result in a new function or protein activity (140, 141). Four CYP74 peptides (CsAOS1,2,5 and CsHPL) have been detected in association with *Cannabis* flowers, and of these, CsHPL was the only isoform found within trichome heads and stalks (132).

### AOC gene family

AOC activity yields the first JA and is the last enzymatic step of JA biosynthesis performed in the chloroplast (101). Three members of the AOC gene family were identified in *C. sativa*, a number consistent with four of *Arabidopsis* (33), six of soybean (142), and two of maize (26). Isoforms typically display organ- and tissue-specific expression and function as heterodimers (143). Only CsAOC2 and CsAOC3 were predicted to localize to plastids (Table 1), supporting the function of these isoforms in JA-biosynthesis. Remarkably, CsAOC3 showed synteny with both *Arabidopsis* (*AtAOC1* and *AtAOC4)* and tomato (*SlAOC*) orthologues. In *Arabidopsis*, heteromers containing AtAOC1 and AtAOC4 display the greatest enzymatic activity (144), suggesting an evolutionary advantage of this conserved genomic region for JA production.

### OPR gene family

A larger variability in OPR gene content exists across plant species analyzed so far: three in *Arabidopsis* (38), five in pea (145), 5 in watermelon (146), 8 in maize (147), 10 in cotton (148), 13 in rice (149), and 48 in wheat (150). This study identified one Type I OPR and four Type II OPRs in the *C. sativa* genome. Both subgroups reduce α, β -unsaturated double bonds and are involved in reactive electrophilic species (RES) detoxification (151, 152). While Type II OPRs are well-characterized for their peroxisomal role in reducing 12-OPDA to OPC8:0 during JA biosynthesis, Type I OPRs are less understood. They have been shown to reduce 4, 6-trinitrotoluene (TNT) (93) and were recently demonstrated to be involved in producing JA through reduction of an endogenous cyclopentenone JA-analog (42). Tandem duplication has been implicated as playing a major role in the size of OPR gene families across plant species (153) and interestingly, while all Type II OPRs were found together in a in a gene cluster, no OPR displayed synteny across the species tested, suggesting OPR gene duplication was a relatively recent event in the *C. sativa* ancestor.

### Conclusion

While the LOX pathway is the hypothetical origin of the hexanoic acid moiety used for cannabinoid biosynthesis (12, 19, 24, 25, 96), the precise mechanisms for its production are largely unknown. Here, we provide a comprehensive description of the *C. sativa* LOX pathway, including the major oxylipin biosynthetic gene families, to establish a working model (Fig. 10) for further investigations.

**Fig. 10.**
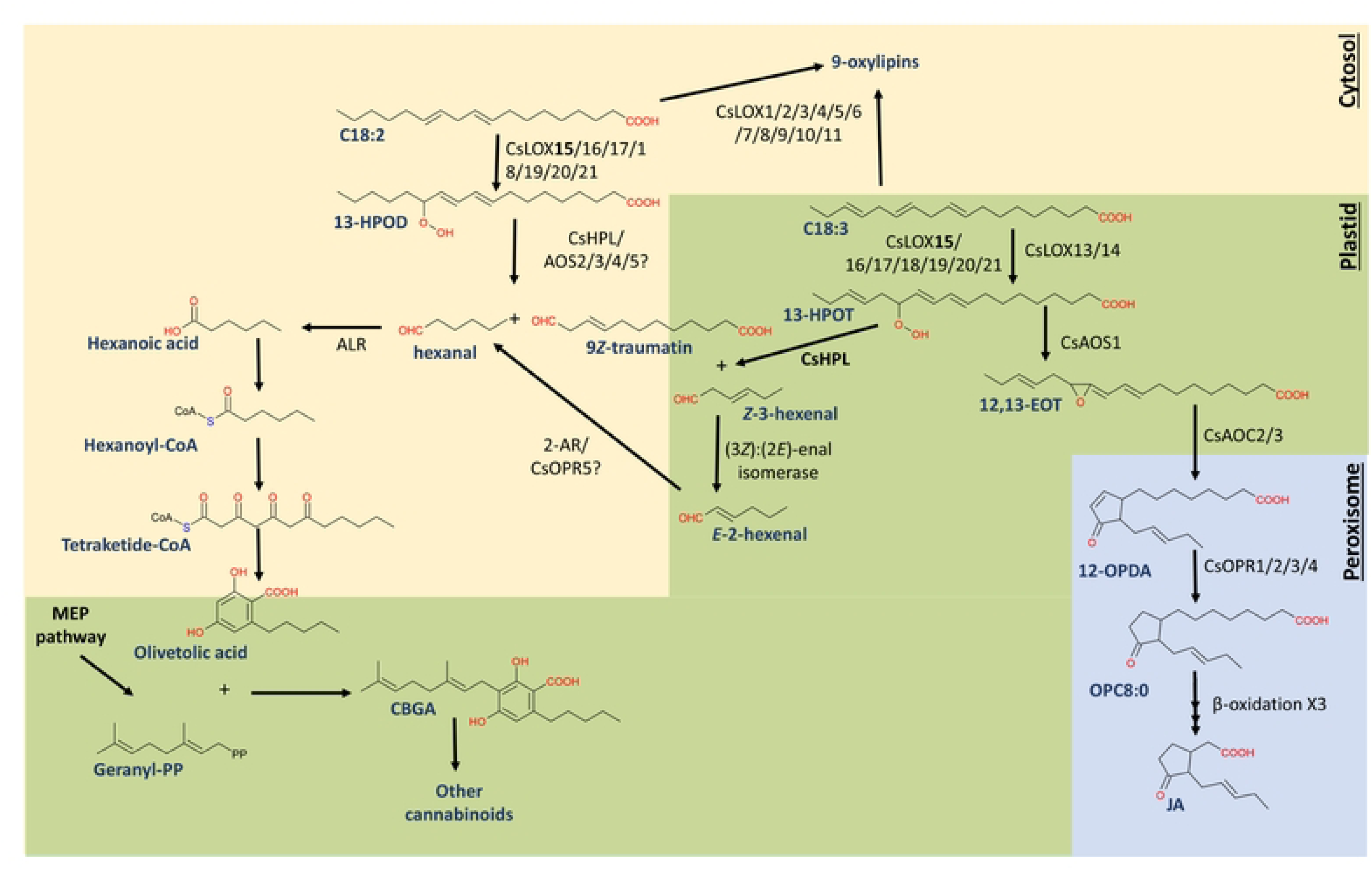
Working model of *C. sativa* oxylipin biosynthetic enzyme isoforms and pathways involved in cannabinoid, GLV, and JA biosynthesis. <u>Abbreviations</u>: [enzymes]: allene oxide cyclase (AOC), allene oxide synthase (AOS), 2-alkynal reductase (2-AR), hydroperoxide lyase (HPL), lipoxygenase (LOX), methylerythritol 4-phosphate pathway (MEP), 12-oxo-phytodienoic reductase (OPR); [metabolites]: (9*Z*, 11*E*, 13*S*, 15*Z*)-12,13-epoxy-9,11,15-octadecatrienoic acid (12,13-EOT), (9*Z*, 11*E*, 13*S*)-13-hydroperoxy-9,11-octadecadienoic acid (13-HPOD), (9*Z*, 11*E*, 13*S*, 15*Z*)-13-hydroperoxy-9,11,15-octadecatrienoic acid (13-HPOT), cannabigerolic acid (CBGA), linoleic acid (C18:2), alpha-linolenic acid (C18:3), jasmonic acid (JA), 12-oxo-phytodienoic acid (12-OPDA), 8-[3-oxo-2-cis-(25)cyclopentyl]octanoic acid (OPC8:0).

Within this context, it will become important to understand the co-localization of the substrates with their corresponding enzymes. LOX activity depends on a 1,4-dipentadiene structure found in either C18:2 or C18:3 (27), both of which, in tomato, account for the most abundant fatty acids in its trichomes (154). However, while HPL-cleavage of 13-hydroperoydienoic acid would generate a statured C_6_-aldehyde, the GLV-producing 13-LOXs and HPL are likely localized to the C18:3-rich chloroplasts (155, 156). Additionally, the GLV-producing AtLOX2 oxygenates C18:3 more efficiently than C18:2 (104), which together may explain why unsaturated GLVs prevail as the major C_6_ volatile in plants (157).

Enzymatic production of hexanoic acid from the C18:3 HPL-product, *Z*-3-hexenal is unexplored, but would reasonably require distinct enzymatic reactions. Chemical or enzymatic isomerization by (3Z):(2E)-enal isomerase would generate (2*E*)-hexenal (46, 55) which could be reduced to hexanal by 2-alkenal reductase (158) or by OPR (159). The conversation of the fatty aldehyde to fatty acid likely requires an aldehyde dehydrogenase (160), the product of which would be readily diffused across cell membranes (161). In an alternative scenario, 13-LOX may oxygenate C18:2 directly, however this would presumably require the cytosolic localization of both 13-LOX and HPL. This strategy would take advantage of the relatively high C18:2 content in the inflorescences of some *Cannabis* cultivars (162).

Understanding and applying knowledge of the *Cannabis* oxylipin biosynthetic pathway is expected to provide novel environmentally friendly approaches towards improving the crop and its desirable consumer traits. Similar technologies in maize and soybeans in development aim at increasing disease resistance (163), drought tolerance (164), and seed flavor (165). Exploiting the diversity in *Cannabis* cultivars via oxylipin biosynthetic gene expression and variability in the predominant oxylipin biosynthetic pathway branch across cultivars and species (e.g., *C. sativa* vs *C. indica*) should also provide opportunities for manipulating cannabinoid content and other traits through marker-assisted selection or identification of superior alleles.

## 5 Conflict of Interest

The authors declare that the research was conducted in the absence of any commercial or financial relationships that could be construed as a potential conflict of interest.

## 6 Author Contributions

EB conceived and designed the study. EB, MR, JT, and EE collected the data. EB, JT, and EE performed analysis. EB wrote the manuscript with input from EE and all authors reviewed and agree with the final version.

## 7 Acknowledgments

Thank you to Drs. Michael Kolomiets and Zachary Schultzhaus for helpful discussions and to Dr. Michael Osier for use of computational resources.

## 8 Funding

We would like to thank the Thomas H. Gosnell School of Life Sciences (GSoLS) and the College of Science (COS) at the Rochester Institute of Technology (RIT) for ongoing support.

## 10 Supporting information

**S1 Fig.**
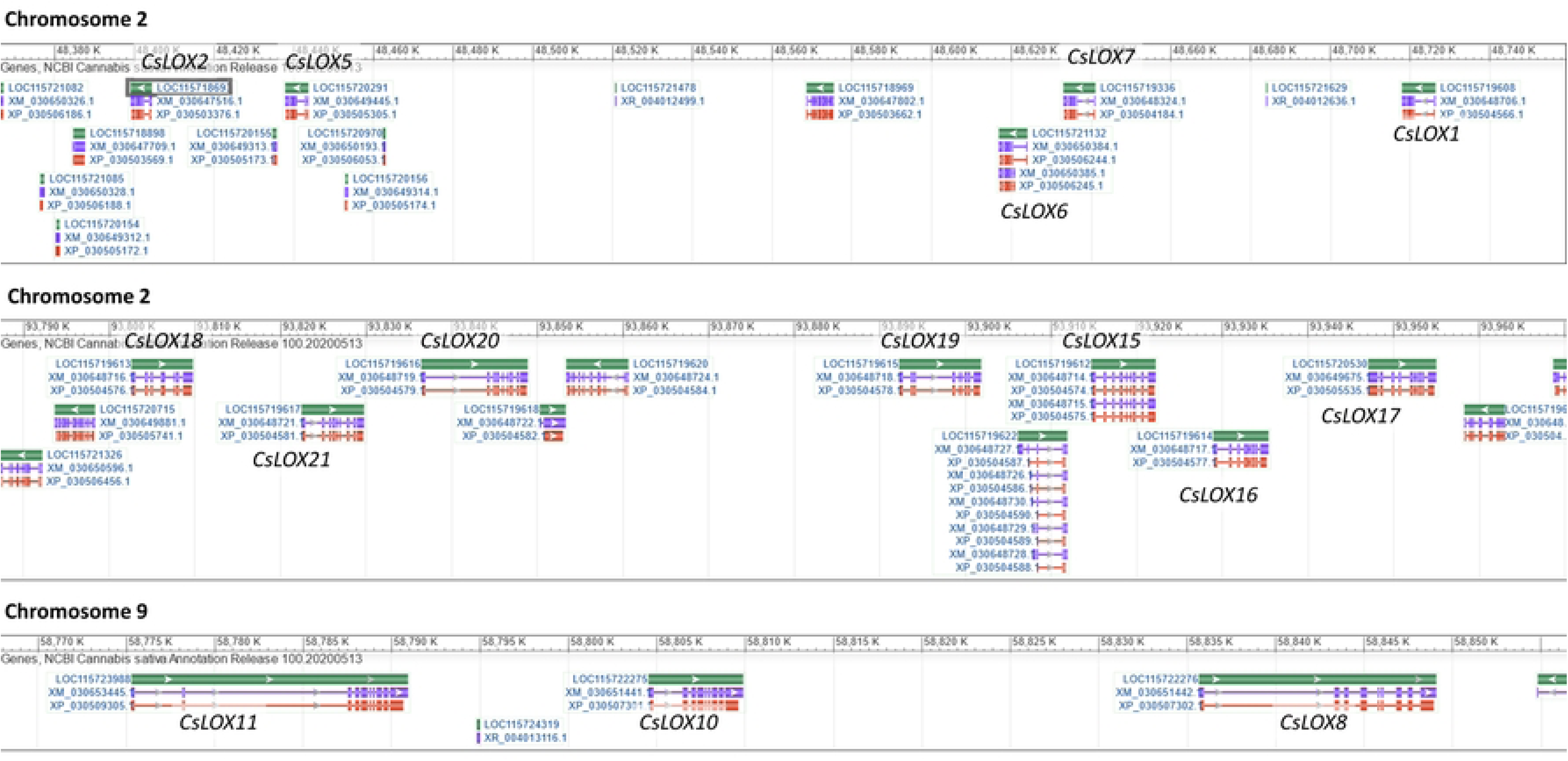
Regions of duplicated LOX genes on *C. sativa* Chromosomes 2 and 9.

**S2 Fig.**
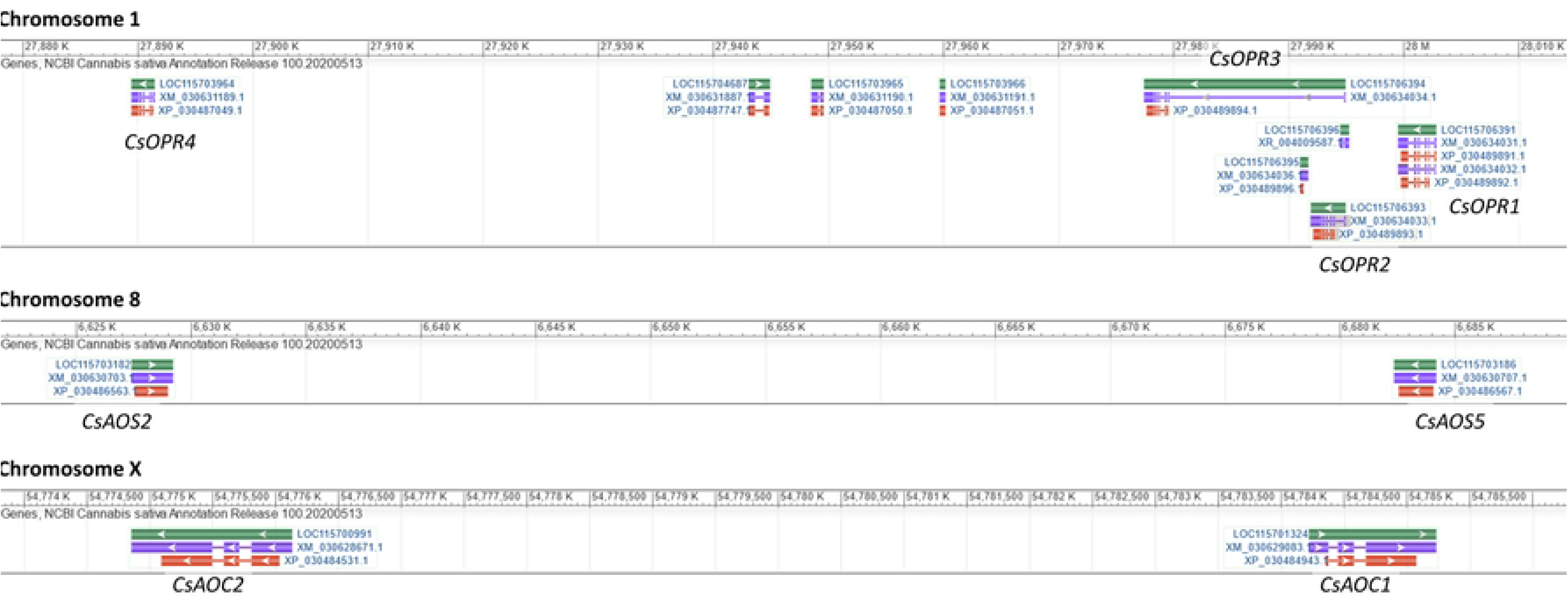
Regions of duplicated *OPR*, *AOS*, and *AOC* genes on *C. sativa* Chromosomes 1, 8, and X.

**S3 Fig.**
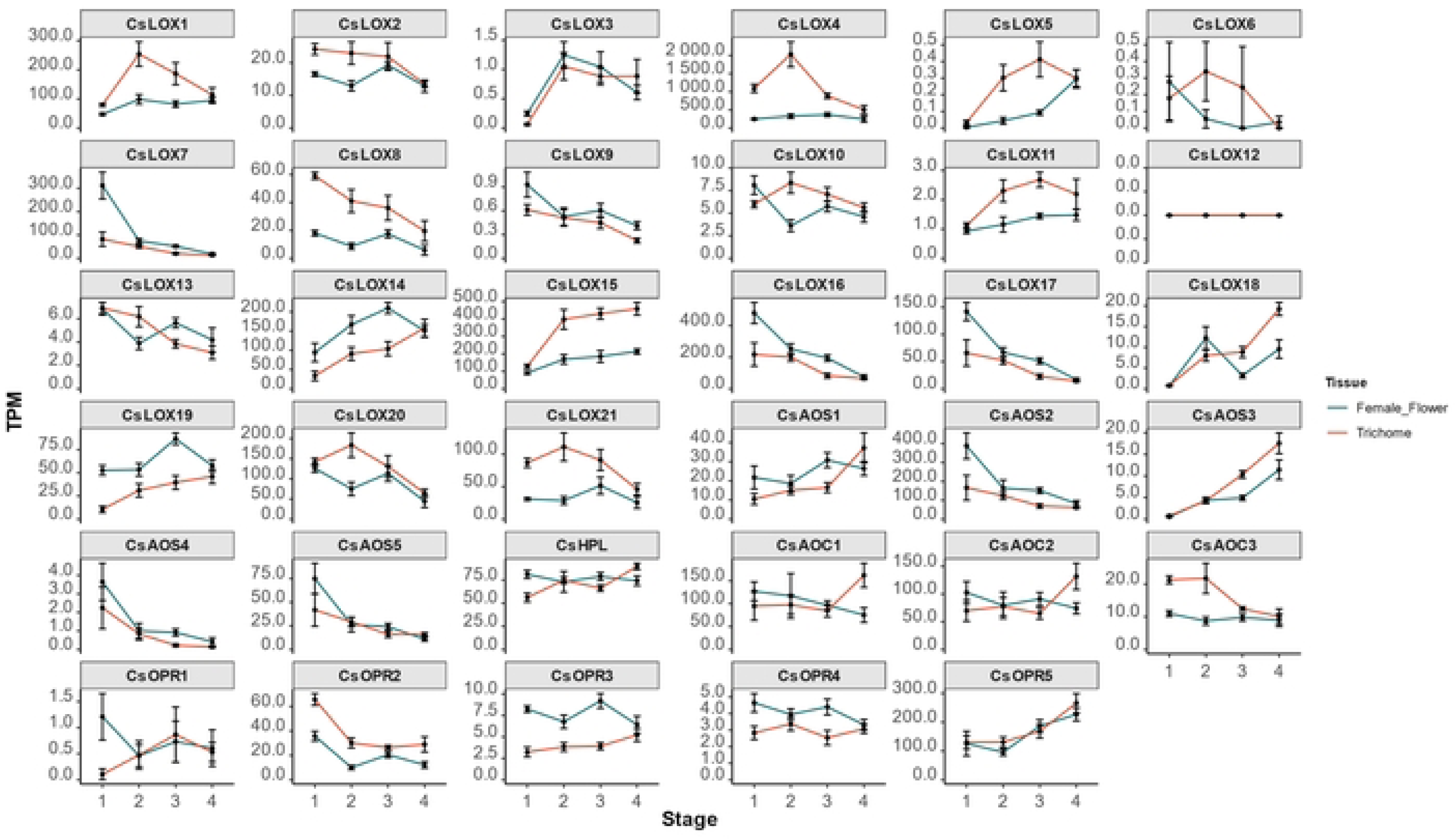
Expression of oxylipin biosynthetic genes across four developmental stages in *C. sativa* flowers and trichomes.

## Notes

### Competing Interest Statement

The authors have declared no competing interest.

